# Impact of Focused Ultrasound on the Cellular Network of Liver Tissue: A New Perspective for Thermal Lesion Detection

**DOI:** 10.1101/2025.10.02.680035

**Authors:** Adrien Rohfritsch, Alexis Griffon, Elorri Olhagaray, Antoine Bienassis, Laura Barrot, Pauline Muleki-Seya, David Melodelima

## Abstract

**Objective:** The noninvasive characterization of soft tissue microstructure remains challenging and has a significant clinical impact on diagnosis and therapy monitoring. During high-intensity focused ultrasound (HIFU) treatments, coagulation necrosis is accompanied by mechanical changes. The objective of this work is to use the anisotropy arising at the cellular level as a new biomarker for treatment evaluation.

**Approach:** We demonstrate that HIFU induces anisotropic alterations in the cellular architecture of liver tissue, which are detectable through the angular dependence of the backscattering coefficient (BSC). Also, in vivo experiments reveal a distinct anisotropic histological pattern localized in the HIFU-treated region.

**Main results:** We show that the degree of anisotropy deduced from BSC measurements is correlated with the histological observations. Moreover, anisotropy increases with delivered energy, providing a quantitative link between treatment parameters and tissue response.

**Significance:** These findings establish BSC anisotropy as a previously unexplored signature of thermal lesions, offering a promising approach for monitoring and feedback in thermal therapeutic ultrasound applications. This breakthrough could open the door to next-generation imaging tools, accelerating the widespread adoption of this highly effective therapeutic modality.

## 1 Introduction

The noninvasive detection of microstructural tissue properties is a key challenge in medical diagnosis and for the monitoring of targeted therapies. The design of novel ultrasound (US) imaging methods has become the focus of extensive research ^1,2^. This modality offers the advantages of low cost and wide accessibility. Moreover, US wave propagation is inherently sensitive to the tissue microstructure.

The characterization of anisotropic structures represents a major topic at the heart of important clinical challenges, such as assessing the myocardial architecture ^3,4^, tumoral microenvironment^5^, renal microstructure ^6^, renal fibrosis ^7^, the orientation of circulating red blood cells ^8^ and muscle characterization ^9,10,11^. Measuring the backscattering coefficient (BSC) at various insonification angles offers a straightforward way to locally estimate microstructure anisotropy, as it is intrinsically dependent on the scatterer organization ^12^. This BSC was previously used by Guerrero ^10^ and Santoso ^11^ to estimate muscle orientation. Furthermore, the use of plane wave imaging (PWI) ^13^ allows for quantifying this anisotropy through precompounded BSC images (obtained for each steering angle) and thus is compatible with real-time imaging ^8^.

High-intensity focused ultrasound (HIFU) alters the tissue microstructure and induces mechanical changes. This noninvasive and nonionizing technique is capable of precisely ablating biological tissues through localized thermal effects. Despite its proven efficacy, its clinical adoption remains limited by the lack of reliable monitoring biomarkers compatible with ultrasound imaging guidance. Thermal necrosis is typically assessed using MRI ^14^, X-ray computed tomography (CT) ^15^, or conventional B-mode ultrasound ^16^. Among these modalities, B-mode imaging is the most accessible but suffers from critical limitations. Notably, changes in echogenicity are often transient and are known to vary significantly across different tissue types ^17^. To overcome these limitations, several quantitative parameters and processing methods have been proposed as ultrasound-based biomarkers for assessing the area ablated by HIFU treatments. Among the most important are tissue elasticity ^18,19^, backscattered energy ^20,21^, Nakagami imaging ^22^, entropy imaging ^23^, harmonic motion imaging ^24^ and parameters derived from the BSC ^25^. While most of these methods rely on the analysis of backscattered radiofrequency (RF) signals, a precise understanding of the underlying microstructural changes remains incomplete. In this context, a recent study reported the changes in the BSC following thermal ablation and linked them to variations in the mean cell diameter ^26^, highlighting the crucial role of the cellular network in the scattering response of human liver tissue. These insights are essential for the development of new imaging tools aimed at assessing the structural changes induced by necrosis.

In this work, we show for the first time that HIFU treatment induces anisotropy at the cellular level in liver tissue. This anisotropy is observed both ex vivo and in vivo through histology and quantified through spectral analysis of binarized cellular network images. Afterward, the first ultrasound measurement of this anisotropy at the local scale based on BSC measurements is reported. These results highlight an unexplored mechanism of HIFU and open up the pathway to exploit it as a new biomarker for HIFU treatment assessment with ultrasound.

## 2 Materials and Methods

### 2.1 HIFU thermal ablation setup

#### 2.1.1 Ex vivo experiment

Three ex vivo liver samples were treated with HIFU at various intensities. Liver samples were purchased from a local slaughterhouse. Before the experiment, the tissues were degassed for 30 minutes, placed on an acoustic absorber in a tank filled with degassed water, and heated to 37°C. The treatment device features an air-backed toroidal HIFU transducer designed for use as a volumetric tissue ablation device in the laboratory. It is composed of 24 piezoelectric transducers emitting at 1.4 MHz, enabling large thermal ablations without mechanical movement of the probe. An imaging probe integrated into the center of the therapy probe allowed visualization of the focal zone during sonication.

These power values are used to estimate the amount of energy deposited at the focus, *E*_*F*_, which is given by (1)

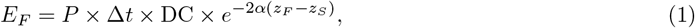

where *z*_*F*_ and *z*_*S*_ represent the focal depth and the surface of the liver, respectively, with respect to the surface of the HIFU probe. In this work, we set *α* = 0.61dB/(cm. MHz), in agreement with values from the literature ^26,27^.

Table 1 summarizes the parameters used to create thermal ablations in the liver samples.

**Table 1:** Summary of the different samples presented and the thermal treatment conditions applied.

| Sample | Treatment | Treatment power ($P$) | Treatment duration ($\Delta t$) | Duty cycle (DC) | $E_F$ |
| --- | --- | --- | --- | --- | --- |
| 1 (ex vivo) | HIFU | 60W | 30s | 100% | 1.3 kJ |
| 2 (ex vivo) | HIFU | 60W | 45s | 100% | 2.0 kJ |
| 3 (ex vivo) | HIFU | 80W | 45s | 100% | 2.7 kJ |
| 4 (ex vivo) | Water bath heating | 70°C | 600s | ∅ | ∅ |
| 5 (in vivo) | HIFU | 21W | 160s | 80% | 2.7 kJ |
| 6 (in vivo) | HIFU | 16W | 160s | 80% | 1.9 kJ |

#### 2.1.2 In vivo experiment

In accordance with European directive 2010/63/EU, all animal procedures were reviewed and approved by the ethical committee n°010 and authorized by the Ministry of Higher Education and Research (#50384-2024062415341673 v4). The protocol was initially developed for a separate scientific objective, and the results presented here were analyzed in parallel to those of the primary study. The HIFU probe and experimental setup were the same as those described in ^21^.

The experiments were conducted on three Landrace pigs, 10-18 weeks old, weighing between 37 and 38 kg. A porcine model was chosen because its anatomy and physiology are similar to those of humans. The animals were kept on site 7 days before the start of the experiments and were fasted 24 h before the US procedure. Premedication was achieved using an intramuscular injection of a mixture of ketamine 1000 (Virbac France, 10-12 mg/kg^−1^), StresnilÂ® (Lilly France, 4-5 mg/kg^−1^) and atropine (Renaudin France, 1 mg) 15 minutes before anesthesia. A 20-gauge catheter was then placed in an auricular vein. Propofol (Fresenius Kabi France) was used to induce (2 mg/kg) and maintain (3 mg/kg/h) anesthesia. During the entire procedure, a continuous infusion (5 *µ*g/kg/h) of sufentanil (Janssen-Cilag France) was administered. The animal was intubated (6 mm endotracheal tube) and placed under assisted ventilation. Oxygenation was supplied by an assisted ventilation system (Vivo 65, Breas Medical, Billerica, MA, USA) at a rate of 7.2 L/min and a frequency of 12 cycles/min (duty cycle: 40%). Abdominal surgery was performed, which consisted of a 25 cm wide laparotomy. A section protector-retractor Alexis was placed, and the liver lobes were exposed. The HIFU probe was placed in contact with the liver parenchyma.

The HIFU treatment consisted of 16 cycles of 10 seconds each, with a duty cycle of 80%. Two in vivo HIFU lesions were obtained with an acoustic power of 21 W and 16 W in water (the corresponding *E*_*F*_ values are reported in Table 1). At the end of the experiment, the anesthetized animal was euthanized using 10 mL of EuthoxinÂ® (Chanelle Pharmaceuticals Manufacturing Irlande, 114 mg/kg). HIFU lesions were sampled and fixed in formaldehyde and then histologically analyzed using the same protocol as that used for the ex vivo samples.

### 2.2 Water bath heating procedure

To better understand the specific effects of HIFU-induced thermal treatment, a liver tissue sample was heated in a water bath. After being degassed for 30 minutes, the sample was placed in a tank filled with degassed water heated to 70°C. A magnetic stirrer ensured a homogeneous temperature distribution in the tank. A needle thermocouple was inserted into the tissue to monitor the internal temperature. After 10 minutes, the sample reached thermal equilibrium with the surrounding water and was immediately transferred to a tank of degassed water at 37°C for subsequent ultrasound imaging and histological analysis.

### 2.3 Histological analysis framework

The tissue samples were fixed, embedded in paraffin blocks, sectioned, and stained with hematoxylin and eosin (H&E) to highlight the cellular architecture. The resulting slides were fully scanned using Zen software (Zeiss, Jena, Germany), enabling detailed, localized analysis of cellular organization across different regions. To characterize the spatial organization of the cellular network within the liver, the tissue was modeled as a biphasic medium, described using a continuous index *I*(**x**) of the form (2):

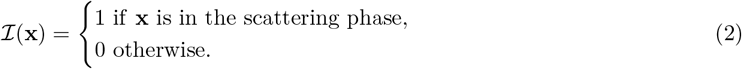

The index ℐ describes the arrangement of the scattering phase (the cellular network) within the host medium (the extracellular matrix). For the sake of understanding, examples of media with increasing degrees of anisotropy are generated numerically and presented in Fig. 1(a), (c) and (e).

**Figure 1:**
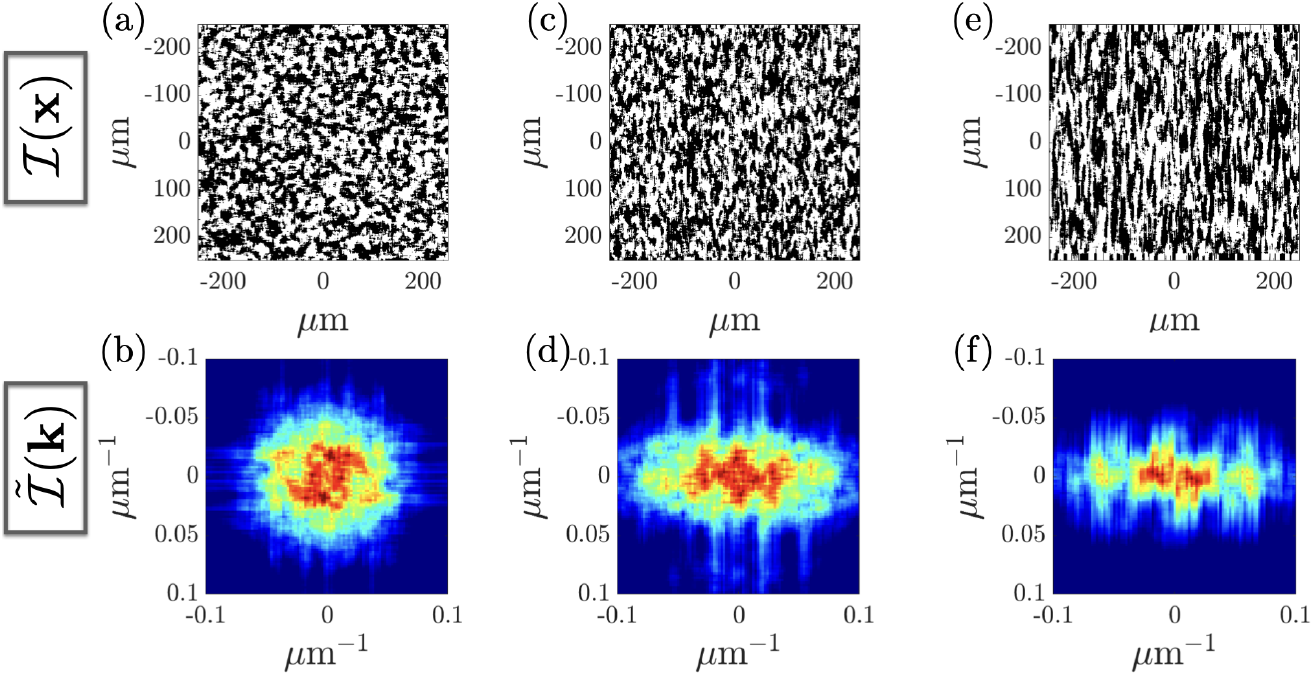
(a), (c), (e): Examples of biphasic continuous media, generated numerically and characterized by the continuous index *I*(**x**), with increasing degrees of anisotropy. (b), (d), and (f) Spectral power densities 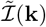 of (a), (c), and (e), respectively.

Determining diffusion from the histological analysis of soft tissue has already been considered in the past. For example, in the work of Mamou ^28^, each medium was assigned an impedance value. Spectral analysis of the resulting impedance map allowed local estimation of the BSC. However, this approach is limited because the impedance values for each phase are often difficult to determine and may be arbitrarily assigned. In this study, we focus instead on local isotropy and anisotropy properties, which are independent of the specific impedance values of each phase. To this end, we analyze the power spectral density of the index ℐ (**x**), denoted 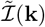 (Fig. 1(b), (d), (f)). This spectral quantity provides information on the angular dependency of the microstructure, as suggested by Zachary ^29^, and therefore also on the angular dependance of the BSC inside this microstructure. The spatial wavevector **k** and the ultrasonic frequency *f* = *c*_0_*k*_0_*/*2*π* can be related through the following expression:

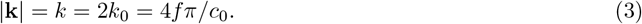

In this work, a constant velocity *c*_0_ = 1540 m/s is assumed for all the samples. All the frequency bandwidths [*f*_min_ − *f*_max_] used to analyze the histological sections presented in this paper are defined on the basis of this assumption.

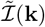 is computed locally over the entire histological section using square regions of 500*µ*m^2^ with a 50% overlap. This region size, corresponding to approximately two wavelengths for the imaging probe used, was chosen as a compromise between the characteristic length of the cellular network on one side and the resolution of the final map on the other side.

To map the degree of anisotropy across the entire histological slide, the angular dependence of 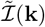 averaged over a given bandwidth is calculated. Noting *k*_*min*_ (*k*_*max*_) the minimal (maximal) value of the spectral interval, the quantity Γ is defined as follows:

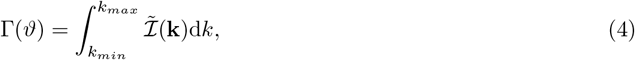

with *ϑ* = arg(**k**).

In the polar plane (*k, ϑ*), if the cellular network is isotropic, Γ(*ϑ*) takes the form of a circle. Conversely, in the case of a strongly anisotropic network, Γ(*ϑ*) takes the form of an elongated ellipse. The ratio between the minor (*d*_1_) and major (*d*_2_) axes of this ellipse is used to define the anisotropy parameter *γ* as follows:

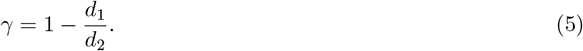

For an isotropic network, *γ* → 0. For an anisotropic medium, *γ* → 1 increases with the degree of anisotropy. This scalar metric therefore quantifies the local anisotropy of the cellular network. The tilt angle of the ellipse, Θ, also provides important information on the microstructure if the microstructure is aligned perpendicular to the HIFU acoustic axis, Θ → 0. Altogether, *γ* and Θ are the parameters that are used in this paper to locally characterize anisotropy.

### 2.4 BSC measurements

The imaging probe integrated into the HIFU device allowed only conventional B-mode imaging. Therefore, an L7-4 probe controlled by a Verasonics ultrasound system was used to image the tissue within the 3.5-7.5 MHz bandwidth. The imaging sequence consisted of seven plane waves steered between ± 18° in 6° increments. Fig. 2 shows numerical simulations of the pressure field profile of the total imaging sequence composed of the seven steered plane wave emissions. RF signals were acquired for each insonification angle, and beamforming was performed using the method described in reference ^30^. It is worth noting that plane wave imaging is not the most conventional imaging method for estimating the BSC, but it has already been the subject of several studies, including the pioneering work of Salles ^13^.

**Figure 2:**
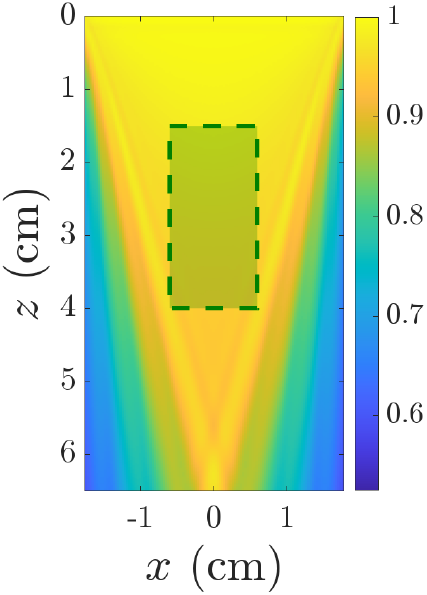
Numerical simulation of the pressure field (normalized amplitude) emitted by the L7-4 probe. The emission sequence consists of 7 plane waves emitted in the *z*-direction, steered by ± 18° (angular step: 6°). The green dotted line rectangle at the center of the field indicates the region where the BSC can be estimated over the full angular aperture.

To estimate the BSC experimentally, one must compensate for the electromechanical system response and the depth-dependent diffraction and focusing effects caused by the ultrasound beam. To achieve this, a common method is adopted here and consists of using a reference phantom, from which the BSC can be determined ^31^. Considering that the RF signals were windowed around the region of interest (ROI) at depth *z* and noting that the power densities inside the sample of interest and inside the reference medium were *P*_meas_ and *P*_ref_, respectively, the measured BSC can be expressed as follows (6):

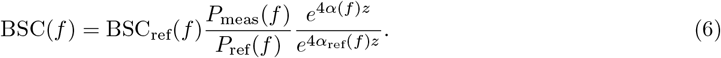

where *α* and *α*_ref_ are the attenuations of the sample and of the reference medium, respectively, accounting for the losses due to absorption and scattering between the interface of the sample and the region of interest. Both quantities were evaluated using the spectral log difference method ^32^ for each sample. The size of the ROI was 20*λ* in both spatial dimensions.

For each insonification angle *θ*, the corresponding spectral power density obtained in the reference phantom under the same insonification direction is used for normalization. The depth of the ROI is also corrected, accounting for the propagation distance at each angle. This leads to the following expression (7):

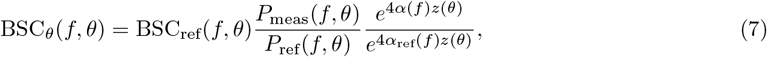

with 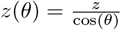. The quantity BSC_*θ*_(*f, θ*) is denoted BSC_*θ*_ when averaged over a frequency range.

The in-house tissue-mimicking phantom used as a reference phantom consisted of polyamide microspheres with a diameter of 5 *±* 2 *µ*m (Orgasol 2001 EXD NAT1, Arkema, France) immersed in agar-agar gel at a concentration of *ϕ* = 1%. The analytic expression of the BSC of this reference medium, BSC_ref_(*f*), is obtained by the so-called “monodisperse structure factor model (SFM)” ^12^ and is given as follows (8):

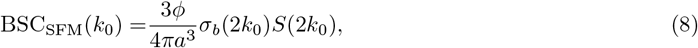

where *a* is the mean radius of the spheres and *σ*_*b*_ is the backscattering cross-section of a single particle, described by Faran’s model ^33^. For a medium made of packings of spherical particles with known density, the structure factor *S* is estimated analytically ^34,35^.

The strategy proposed in this paper involves confronting the dependency of 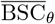 on the insonification angle *θ* and the structural anisotropy studied through histology. An anisotropy mapping metric is also proposed on the basis of the computation of the quantity 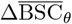, which is defined as follows: (9)

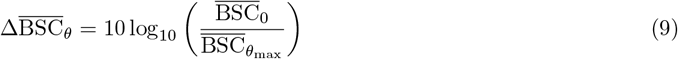

This comparison allows monitoring with ultrasound the emergence of anisotropy during a thermal procedure performed with HIFU.

### 2.5 Fiber angle estimation in anisotropic soft tissue

To validate the ability of the method to measure the orientation of anisotropic microstructures, a porcine tenderloin sample was imaged. This highly fibrous tissue had previously been used by Papadacci to validate their method on the basis of the coherence of backscattered signals ^3^. The advantage of choosing this tissue as a reference medium lies in the fact that the direction of the fibers is visible on the B-mode image. The RF signals were measured with the probe aligned parallel to the fibers. Three positions were selected, and BSC_*θ*_ was estimated for each position within a region of interest (ROI) containing visible fibers.

## 3 Results

### 3.1 Histological quantification of HIFU-induced anisotropy

The presence of HIFU-induced anisotropy is described in Fig. 3. In (a), a full histological section of a liver sample treated *in vivo* with HIFU is presented. In (b) and (f), two smaller sections respectively extracted from the untreated and HIFU-ablated regions of the tissue are shown.

**Figure 3:**
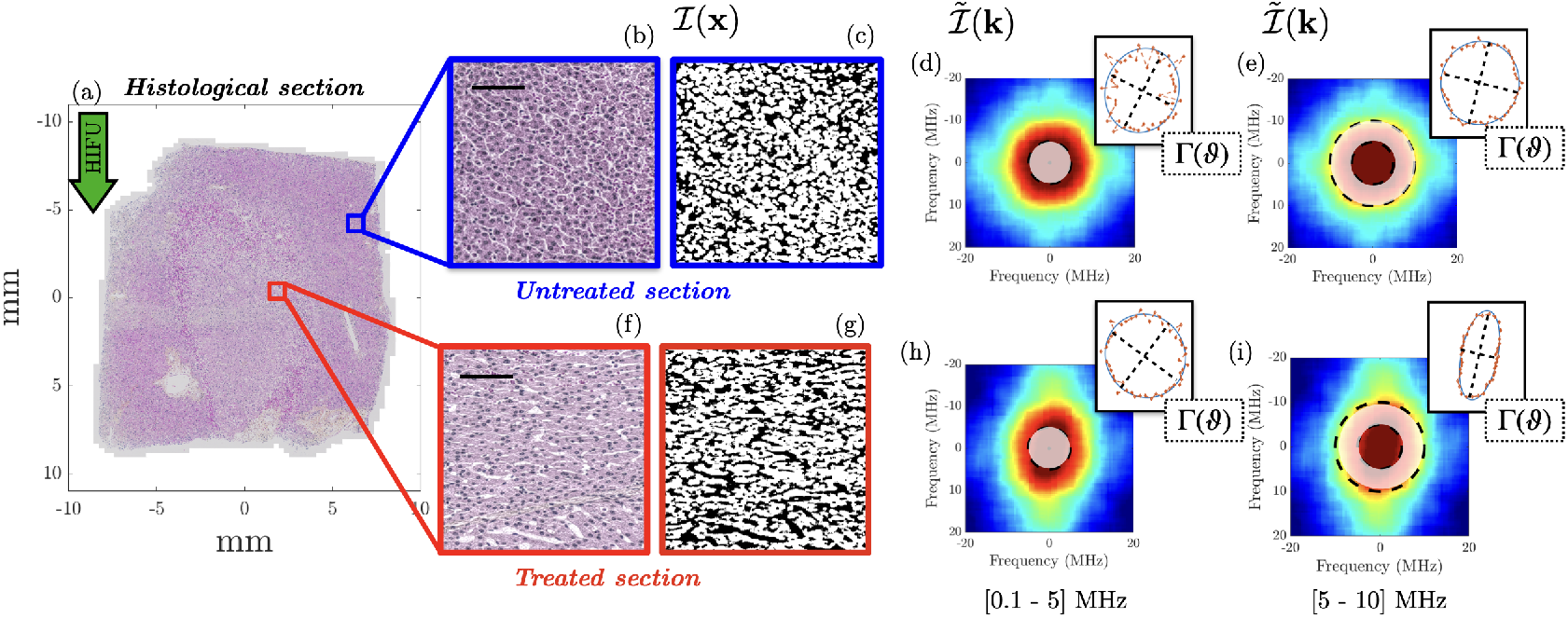
Histological analysis framework. (a) Histological section (staining method: H&E) of an in vivo liver tissue sample treated with HIFU. (b) Zoomed-out view of the untreated area. (c) Binarized version of (b), showing the cellular network in white (scale bar: 100*µ*m). (d, e) Power spectral density 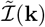 calculated from (c) and polar representation of Γ(*ϑ*) computed for the frequency window [0.1-5] MHz (d) and [5-10] MHz (e). (f-i) Correspond to the same analyses as in (b-e) for a HIFU-treated section of the tissue.

First, the local analysis of the two regions reveals differences in the organization of the cellular network. In the untreated tissue (Fig. 3(b), (c)), the cellular network appears isotropically distributed in the subsection. In contrast, in the treated tissue (Fig. 3(f), (g)), the cellular network exhibits a noticeable alignment perpendicular to the HIFU beam, leading to an anisotropic network.

Analysis of the two microstructures over the [0.1-5] MHz bandwidth reveals no clear evidence of anisotropy in Γ(*ϑ*) (Fig. 3(d) and (h)), and the anisotropy parameter *γ* remains low, with values of 0.09 for the untreated tissue and 0.18 for the treated tissue. In contrast, a marked difference is observed between the two samples when the spectral power densities over the [5-10] MHz bandwidth are analyzed; in the treated section, both the spectral power density 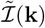 and Γ(*ϑ*) exhibit pronounced angular variations (Fig. 3(i)). While anisotropy remains low in the untreated tissue (*γ* = 0.18) over this frequency range, it increases substantially in the treated sample, reaching *γ* = 0.78 (+333%). Having presented the local characterization of HIFU-induced anisotropy, particular attention is now given to the analysis of the anisotropy parameters *γ* and Θ across entire histological sections of various treatment conditions.

Fig. 4 presents the histological sections of three ex vivo samples: an untreated sample, a sample treated with HIFU above the boiling threshold, and a sample heated to 70°C in a water bath. *γ* and Θ parameter maps were generated for the frequency bandwidth [4-8] MHz, which overlaps with the bandwidth of the ultrasonic probe used for imaging and is relevant for detecting anisotropy. All *γ* and Θ values are reported in Tables 2 and 3. In the case of the HIFU-treated sample, values outside and inside the ablated tissue are reported.

**Table 2:** *γ* values measured on the three ex vivo liver samples presented in Fig. 4.

| Tissue | Figure | Freq. window | $\gamma$ - ablated region | $\gamma$ - untreated tissue |
| --- | --- | --- | --- | --- |
| Untreated liver | 4(a,b) | [4-8] MHz | ∅ | $0.36 \pm 0.14$ |
| HIFU ablated liver | 4(d,e) | [4-8] MHz | $0.40 \pm 0.14$ | $0.32 \pm 0.13$ |
| Water bath heated (Sample 4) | 4(g,h) | [4-8] MHz | $0.34 \pm 0.14$ | ∅ |

**Table 3:** |Θ| values measured on the three ex vivo liver samples presented in Fig. 4.

| Tissue | Figure | Freq. window | $\vert \Theta\vert $ - ablated region | $\vert \Theta\vert $ - untreated tissue |
| --- | --- | --- | --- | --- |
| Untreated liver | 4(a,c) | [4-8] MHz | ∅ | $19 \pm 12$ |
| HIFU ablated liver | 4(d,f) | [4-8] MHz | $9.6 \pm 7.2$ | $17 \pm 12$ |
| Water bath heated (Sample 4) | 4(g,i) | [4-8] MHz | $23.5 \pm 13$ | ∅ |

**Figure 4:**
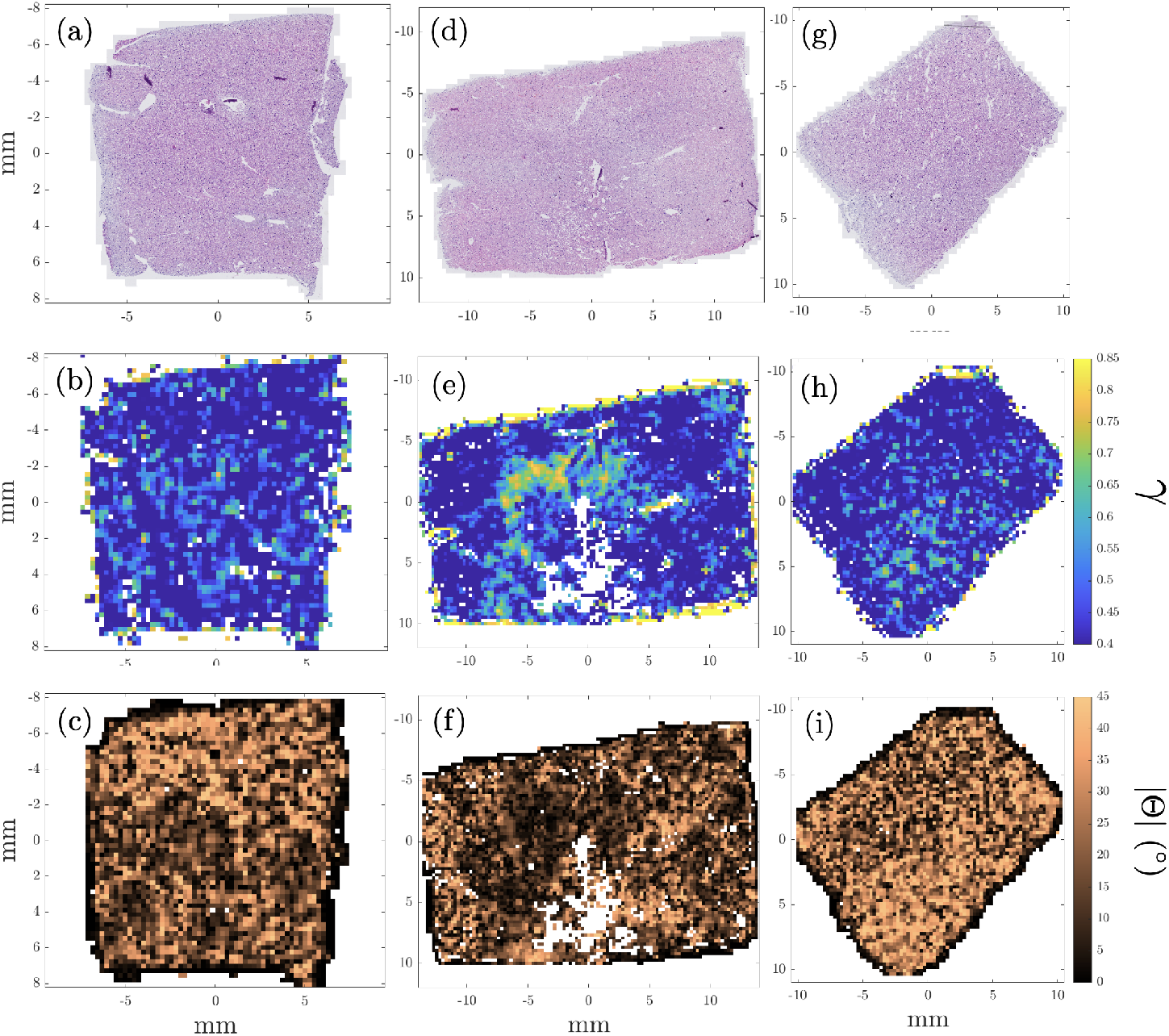
Histological sections (staining method: H&E) of (a) an untreated sample, (d) an ex vivo sample treated by HIFU and (g) a sample heated in a water bath. (b), (e) and (h) Maps of parameters *γ* obtained from (a), (d) and (g), respectively, for a bandwidth of [4 − −8] MHz;. (c), (f) and (i) Maps of parameters |Θ| obtained from (a), (d) and (g), respectively, for a bandwidth of [4 − 8] MHz.

The results for the untreated tissue are presented in maps 4(a), (b), and (c). The *γ* and Θ maps ((b), (c)) do not show significant heterogeneity, which is consistent with the relatively homogeneous nature of the hepatocellular structure.

The two other samples are analyzed after thermal ablation. The tissue treated with HIFU (Fig. 4(d), (e), and (f)) shows strong evidence of anisotropy in both parameters. An increase of 25% in *γ* is observed between the treated and untreated regions (Fig. 4(e)) outside the boiling region. The Θ map (Fig. 4(f)) is also highly informative. In the treated region, the Θ values are less heterogeneous than they are outside of the treated region. Furthermore, a clear decrease in Θ is measured in this area, indicating that the cellular matrix is more organized and tends to align perpendicular to the HIFU acoustic axis.

There is no evidence of anisotropy in the analysis of the tissue heated in a water bath at 70°C (4(g), (h), and (i)), either in *γ* or in Θ. Compared with the unheated sample, a difference of approximately 12% is measured in the Θ values, which is close to the standard deviation within the samples, and only a 5.9% difference is measured in *γ*. These findings demonstrate that water bath heating does not induce anisotropy in the hepatic tissue microstructure.

To go deeper into this analysis, two HIFU lesions induced in vivo in porcine liver using two different delivered energies (Fig. 5) are analyzed on two frequency bandwidths: [0.1-5] MHz and [5-10] MHz. This allows us to determine the influence of the frequency window on the detection of anisotropy, which is important when histological findings are compared with ultrasonic measurements. All *γ* values are reported in Table 4 and the Θ values are reported in Table 5.

**Table 4:**
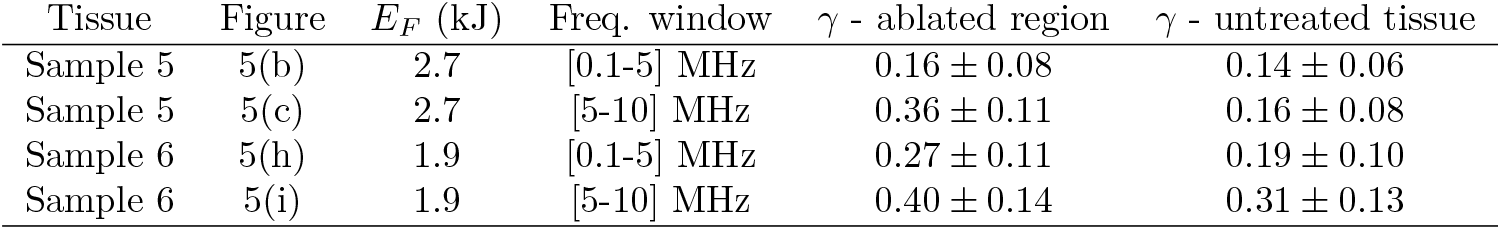
*γ* values measured on the two in vivo liver samples presented in Fig. 5.

| Tissue | Figure | $E_F$ (kJ) | Freq. window | $\gamma$ - ablated region | $\gamma$ - untreated tissue |
| --- | --- | --- | --- | --- | --- |
| Sample 5 | 5(b) | 2.7 | [0.1-5] MHz | $0.16 \pm 0.08$ | $0.14 \pm 0.06$ |
| Sample 5 | 5(c) | 2.7 | [5-10] MHz | $0.36 \pm 0.11$ | $0.16 \pm 0.08$ |
| Sample 6 | 5(h) | 1.9 | [0.1-5] MHz | $0.27 \pm 0.11$ | $0.19 \pm 0.10$ |
| Sample 6 | 5(i) | 1.9 | [5-10] MHz | $0.40 \pm 0.14$ | $0.31 \pm 0.13$ |

**Table 5:** |Θ| values measured on the two in vivo liver samples presented in Fig. 5.

| Tissue | Figure | $E_F$ (kJ) | Freq. window | $\vert \Theta\vert $ - ablated region | $\vert \Theta\vert $ - untreated tissue |
| --- | --- | --- | --- | --- | --- |
| Sample 5 | 5(d) | 2.7 | [0.1-5] MHz | $14.7 \pm 11.2$ | $22.7 \pm 12.3$ |
| Sample 5 | 5(e) | 2.7 | [5-10] MHz | $7.9 \pm 6.5$ | $21.6 \pm 13.0$ |
| Sample 6 | 5(j) | 1.9 | [0.1-5] MHz | $17.3 \pm 12.5$ | $23.6 \pm 13.2$ |
| Sample 6 | 5(k) | 1.9 | [5-10] MHz | $9.7 \pm 8.0$ | $22.3 \pm 13$ |

**Figure 5:**
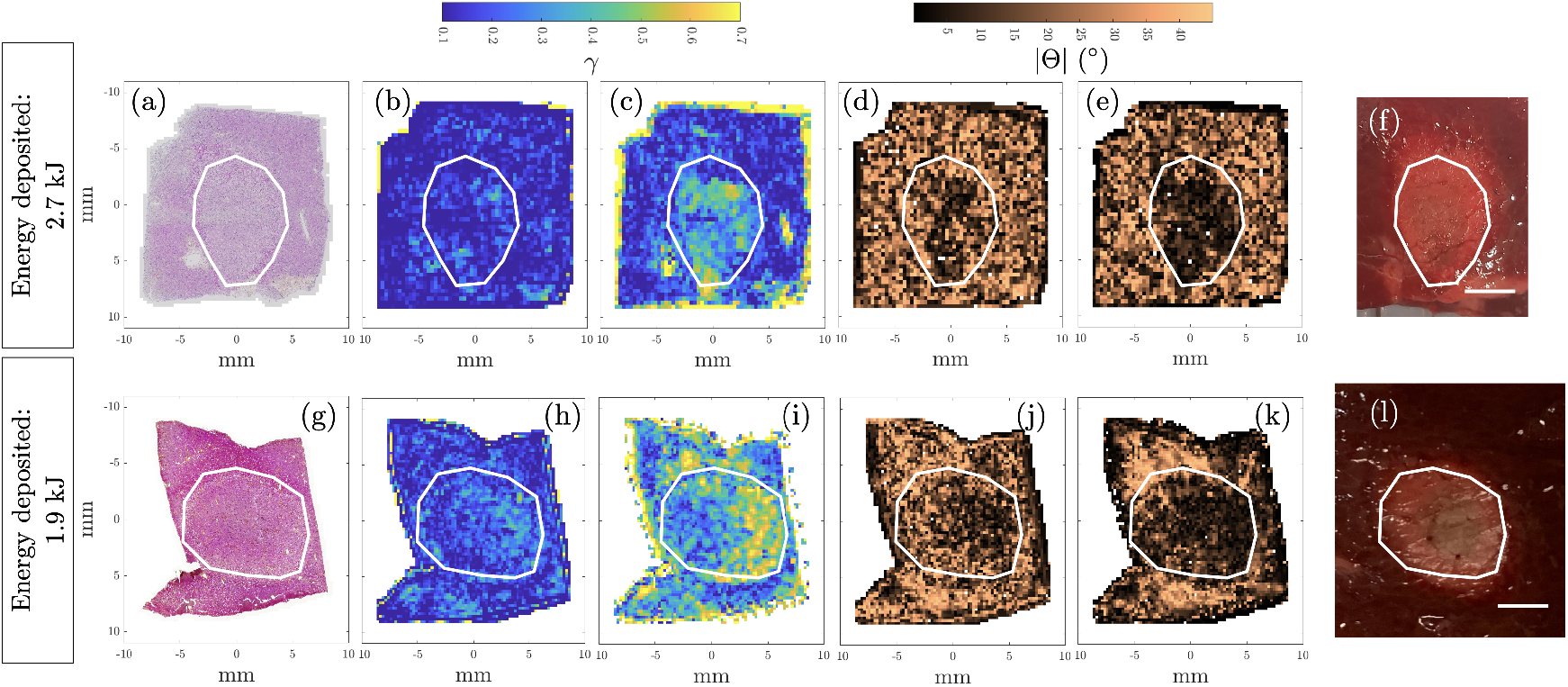
(a), (g) Histological sections (staining method: H&E) of in vivo liver samples treated with HIFU. (b) (respectively (h)) Map of parameter *γ* computed from (a) (respectively (e)) for a bandwidth between [0.1 − 5]MHz. (c), (i) Same as (b), (h) for a bandwidth between [5 − 10]MHz. (d), (e), (j), (k) Represent | Θ |maps corresponding respectively to (b), (c), (h), (i). (f), (l) Macroscopic view from lesions (a) and (g), respectively (scale bar: 5 mm).

In Fig. 5 (a) and (g), histological sections of the HIFU lesions are presented. As shown in Fig. 5(b) and (h), *γ* remains relatively low (*<* 0.3) when the low-frequency bandwidth is considered. At higher frequencies, the entire HIFU-treated region of the sample treated with 2.7 kJ (sample 5) exhibits a strong anisotropic signature, with an increase of 125% between the treated and untreated zones. The HIFU lesion is clearly visible on the *γ* map (Fig. 5(c)) and is in good agreement with the macroscopic observation (Fig. 5(d)). In the case of the sample treated with 1.9 kJ (sample 6), the cellular matrix outside the treated area already shows marked signs of anisotropy (*γ* = 0.31 ± 0.13), highlighting the significant interindividual variability in the hepatocellular structure. Nevertheless, the anisotropy induced by HIFU treatment remains clearly detectable compared with that in the untreated zone, with an absolute difference of 0.9 ± 0.14 and a relative increase of 29%. This lower increase may be attributed to the lower power used during the treatment.

### 3.2 Fiber orientation imaging

Fig. 6 illustrates the measurement of the fiber orientation within the porcine tenderloin sample. For each of the three positions, a distinct peak is observed around the angle identified on the corresponding B-mode images, indicating a predominantly two-dimensional scattering microstructure. Under normal incidence, the increased coherence of the backscattered signals from aligned scatterers leads to higher backscattered energy than under oblique incidence ^4^. These results are thus consistent with the anisotropic structural organization of the tissue and demonstrate the ability of the method to detect the anisotropic structural organization.

**Figure 6:**
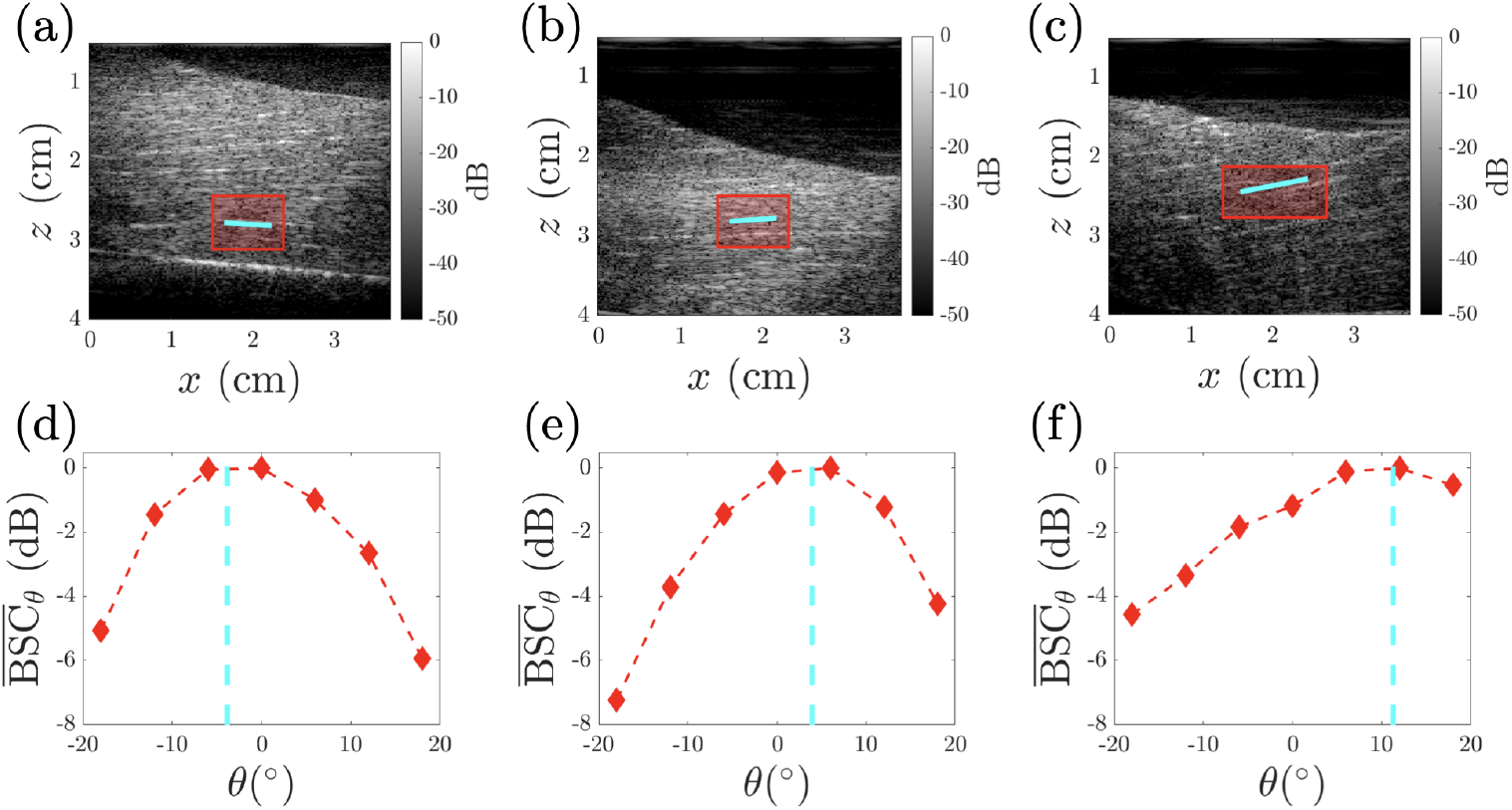
(a), (b), (c): Bmode images at three locations in a porcine tenderloin sample. The red rectangle represents the ROI in which the BSC is calculated, and the cyan line inside indicates the main orientation direction of the fibers within the ROI, measured on the basis of the B-mode image. (d), (e), (f): 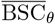 as a function of the insonification angle *θ* obtained from (a), (b), and (c), respectively, and normalized by the maximum value of 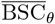 over the angular range ± 18°C. The cyan dotted line indicates the angle measured on the Bmode images.

### 3.3 Imaging of a HIFU-treated sample

In this section, the results obtained by imaging a HIFU-induced lesion created on an ex vivo liver sample and treated at 60 W for 45 seconds (sample 2 in Table 1) are presented. The BSC is first analyzed over two spectral bandwidths: (3.5-4.5) MHz and (6-7) MHz. These frequency bandwidths are selected in accordance with the histological findings reported in the previous section. For each of the seven insonification angles *θ*, BSC_*θ*_ maps are generated. In parallel, spatially averaged BSC_*θ*_ values within and outside the treated region are plotted as a function of *θ*.

Fig. 7 presents the BSC maps for the frequency band (3.5-4.5) MHz. The necrotic region, which is characterized by a localized increase in BSC, is clearly visible in Fig. 7(a). In this case, BSC_*θ*_ does not vary significantly with the insonification angle *θ*. The angular variation in BSC_*θ*_ is at most 1.2 dB within the necrotic region and 1.1 dB in the non-necrotic tissue. This observation is consistent with the histological analysis: at this frequency scale, the cellular network both inside and outside the necrotic area appears isotropic.

**Figure 7:**
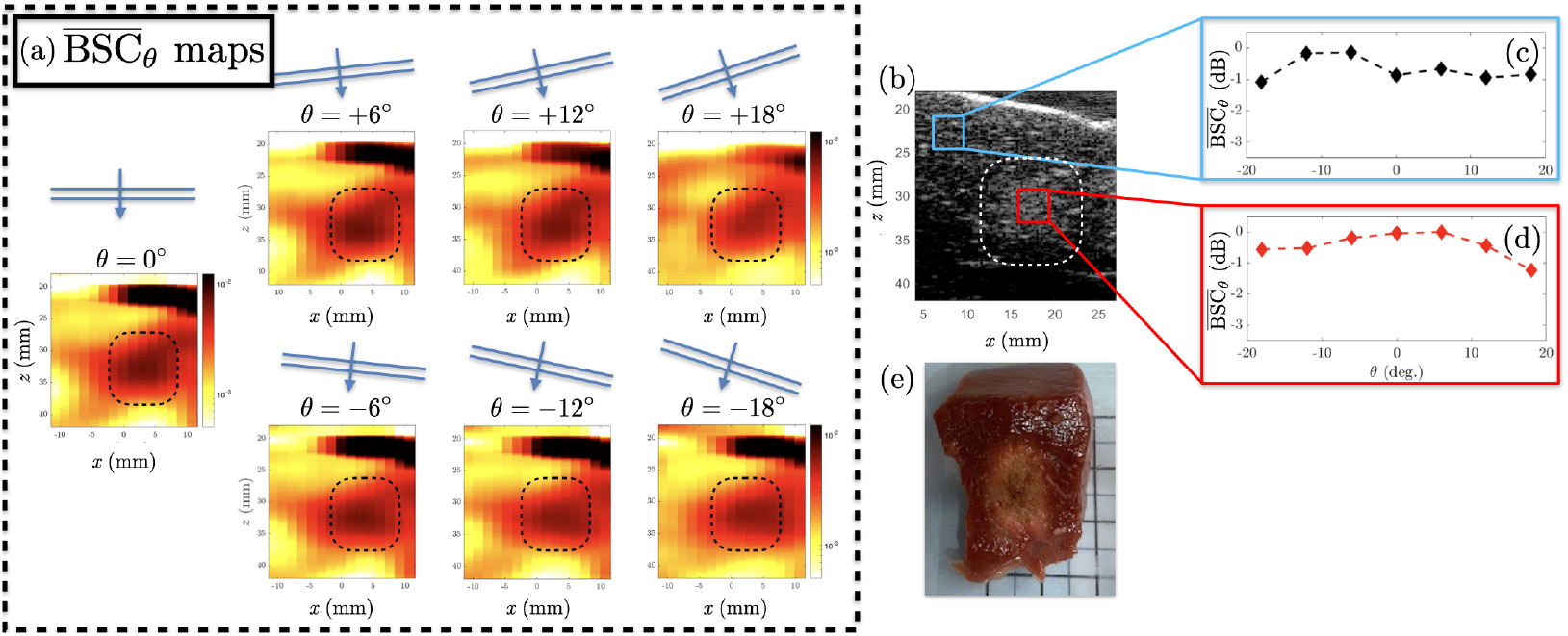
Ex vivo HIFU-treated sample (Sample 2), frequency bandwidth: [3.5-4.5] MHz. (a) 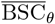 maps obtained for each insonification angle *θ*, inside and outside the HIFU-treated area. (b) B-mode image of the area and BSC value in two ROIs taken outside (c) and inside (d) of the treated area. (e) Macroscopic view of the HIFU lesion. The dotted lines in (a) and (b) qualitatively delineate the ablated region in the tissue.

In contrast, the analysis over the second frequency band (6-7) MHz reveals a different pattern, as shown in Fig. 8. The spatially averaged BSC_*θ*_ within the necrotic region varies significantly with the insonification angle. A difference of 2.8 dB is observed between the normal incidence (*θ* = 0°) and oblique incidence (*θ* = 18°) within the treated area. This anisotropy is not observed outside the necrotic zone, where BSC_*θ*_ varies by less than 1 dB between the two insonification directions.

**Figure 8:**
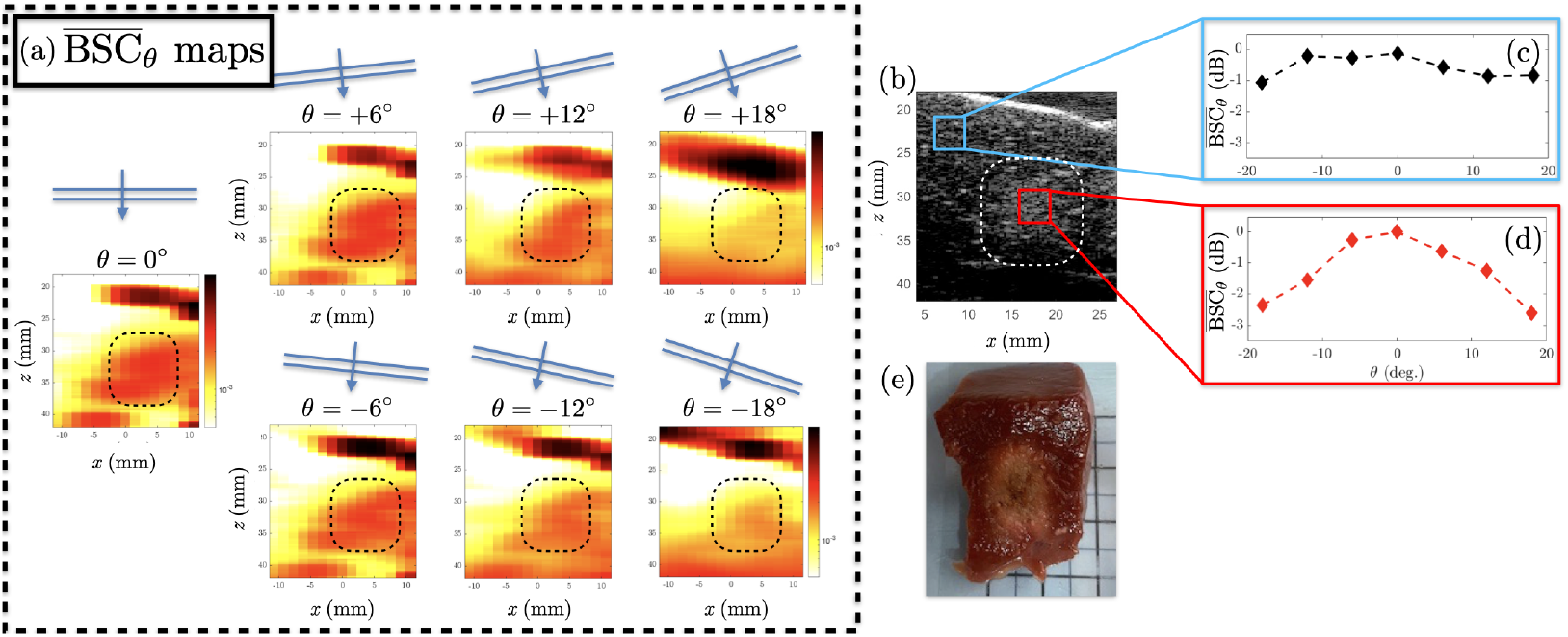
Sample 2 (ex vivo), frequency bandwidth: [6-7] MHz. Same as Fig. 7.

### 3.4 Imaging of a sample heated in a water bath

The results of the BSC measurements conducted on the sample heated in a water bath are presented in Fig. 9. Although the average temperature of 70°C reached in the sample is typical of that achieved during HIFU treatment, no sign of anisotropy appears in the measurement of BSC_*θ*_. Compared with the results of the histological analysis, these findings demonstrate that a heated tissue whose cellular network shows no strong signs of anisotropy is also isotropic from an ultrasonic perspective.

**Figure 9:**
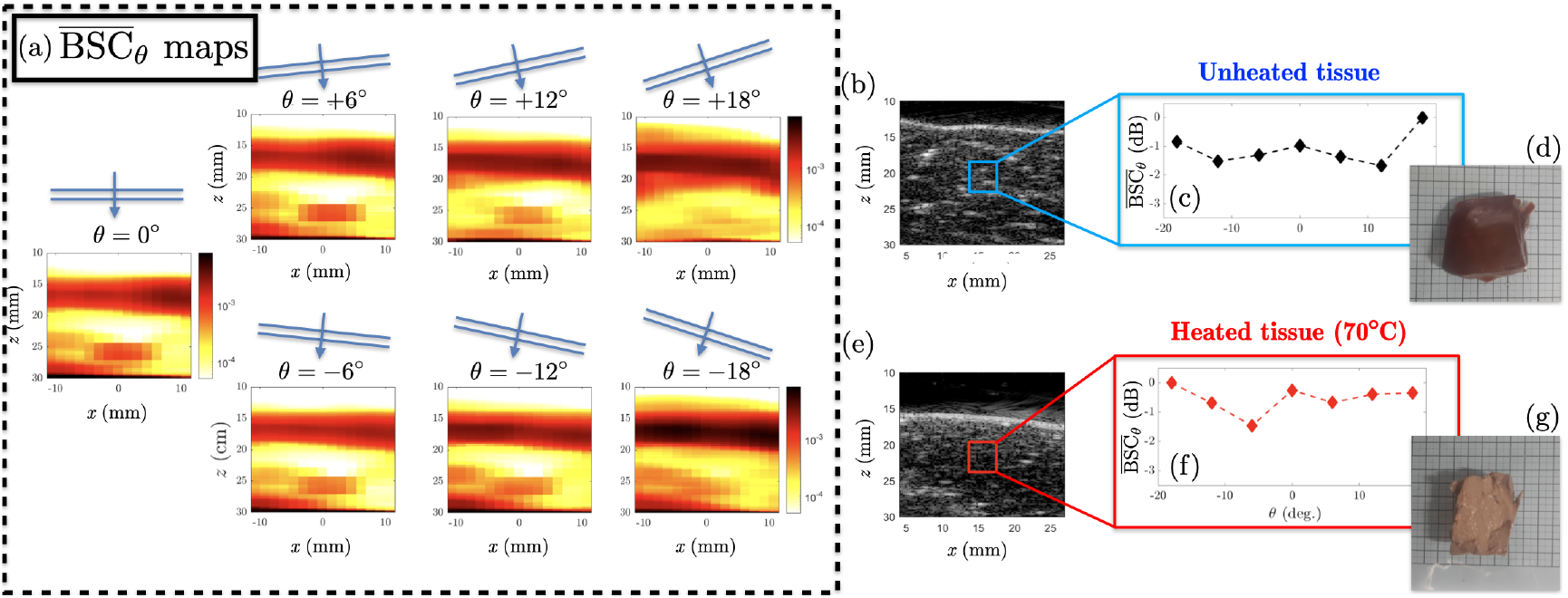
Sample 6 (ex vivo), frequency bandwidth: [6-7] MHz. (a) 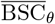 maps of the water bath-heated sample obtained for each insonification angle *θ*. (b) B-mode image of the unheated sample and (c) BSC in an ROI inside the tissue. (e) Macroscopic view of the unheated sample. (f), (g) and (h) are the same as (d), (e) and (f) for the water bath-heated sample, respectively.

### 3.5 Impact of the treatment parameters on anisotropy

In this section, the influence of the treatment conditions on the tissue structural properties and their impact on BSC_*θ*_ are investigated. Fig. 10(a) presents the *γ* parameter averaged over 15 ROIs inside the treated area of samples 1 to 3 treated with energy increases of 1.3 kJ, 2.0 kJ and 2.7 kJ, respectively. The analysis is performed across seven 1-MHz spectral windows spanning 3.5 to 7.5 MHz. In sample 1, *γ* remains stable at approximately 0.45–a relatively low value that is comparable to that of the untreated tissue (see the previous section). In contrast, sample 2 exhibits a strong frequency-dependent variation in *γ*, highlighting the frequency dependance of microstructural anisotropy. These observations are consistent with earlier results (Figs. 5, 7, 8), confirming that both histological and backscatter anisotropies are frequency dependent. Notably, a clear transition from isotropic to anisotropic behavior is observed in sample 2, with *γ* increasing by more than 50% from the lowest to the highest frequency window. Sample 3, subjected to more intense treatment, shows pronounced low-frequency anisotropy and reaches anisotropy levels similar to those of sample 2 at higher frequencies. The overall increase could be explained by an increase in the energy deposited at and around the focus, as discussed in the next section.

**Figure 10:**
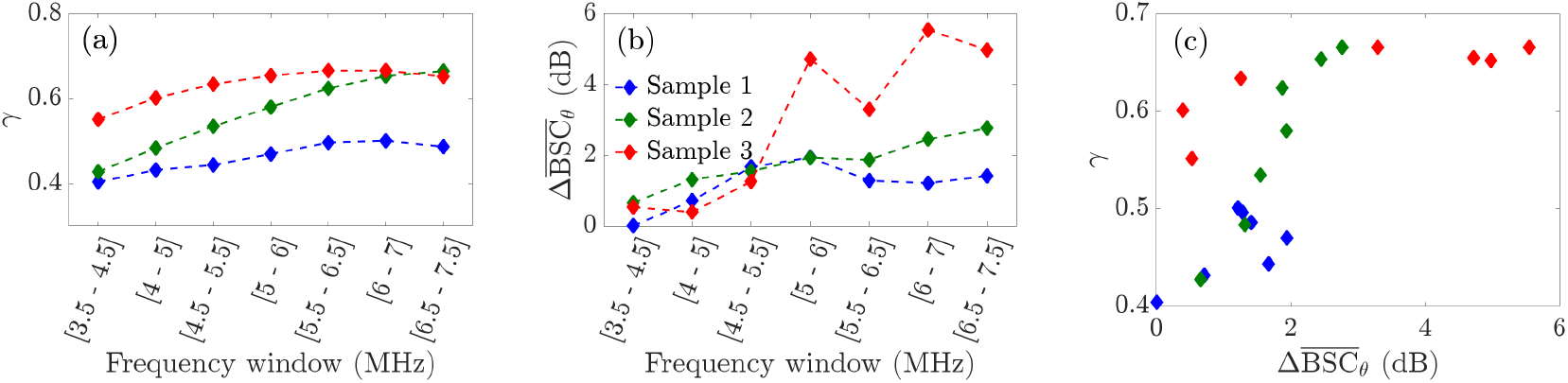
(a) Average value of *γ* as a function of the frequency bandwidth for three ex vivo samples heated with various HIFU sequences. (b) 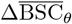 of the same samples, calculated on various frequency bandwidths. (c) Average value of *γ* as a function of 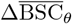.

Fig. 10(b) shows the difference between the minimum and maximum values of 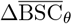 for each spectral bandwidth and for the same three samples. This quantity, denoted as ΔBSC_*θ*_, provides a measure of the degree of anisotropy over a given frequency range. The results are consistent with those in Figure (a): for frequencies above 5 MHz, the energy deposited in the tissue correlates with the level of anisotropy measured with ultrasound. This highlights that anisotropy increases not only with treatment duration (as reflected in the progression from sample 1 to sample 2) but also with higher acoustic power (between sample 2 and sample 3).

The relationship between 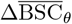 and *γ* is shown in Fig. 10(c). A positive correlation is obtained for samples 1 and 2. In the case of sample 3, a plateau is reached for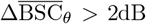. The physical origin of this plateau will be investigated in future work and might be due to other structural alterations that are not contained in the measure of *γ*.

### 3.6 HIFU lesion imaging through anisotropy imaging

To develop a reliable guidance method for identifying treated areas within the imaging plane, a local anisotropy mapping approach is proposed. Here, the selected frequency bandwidth is [6-7] MHz, where strong signs of anisotropy are detected (Fig. 8). Untreated samples and samples 1 to 3 are analyzed. The quantity 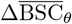 is computed within the region of the imaging plane where all the emitted plane waves overlap. The results are presented in Fig. 11. The absence of anisotropy in the untreated tissue, which is consistent with its isotropic microstructure, is clearly visible in Fig. 11(e). The average value of 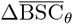 is shown as a function of the energy deposited at the focus in Fig. 11(i). The linear correlation coefficient is *R*^2^ = 0.84, which supports the positive correlation between the two quantities.

**Figure 11:**
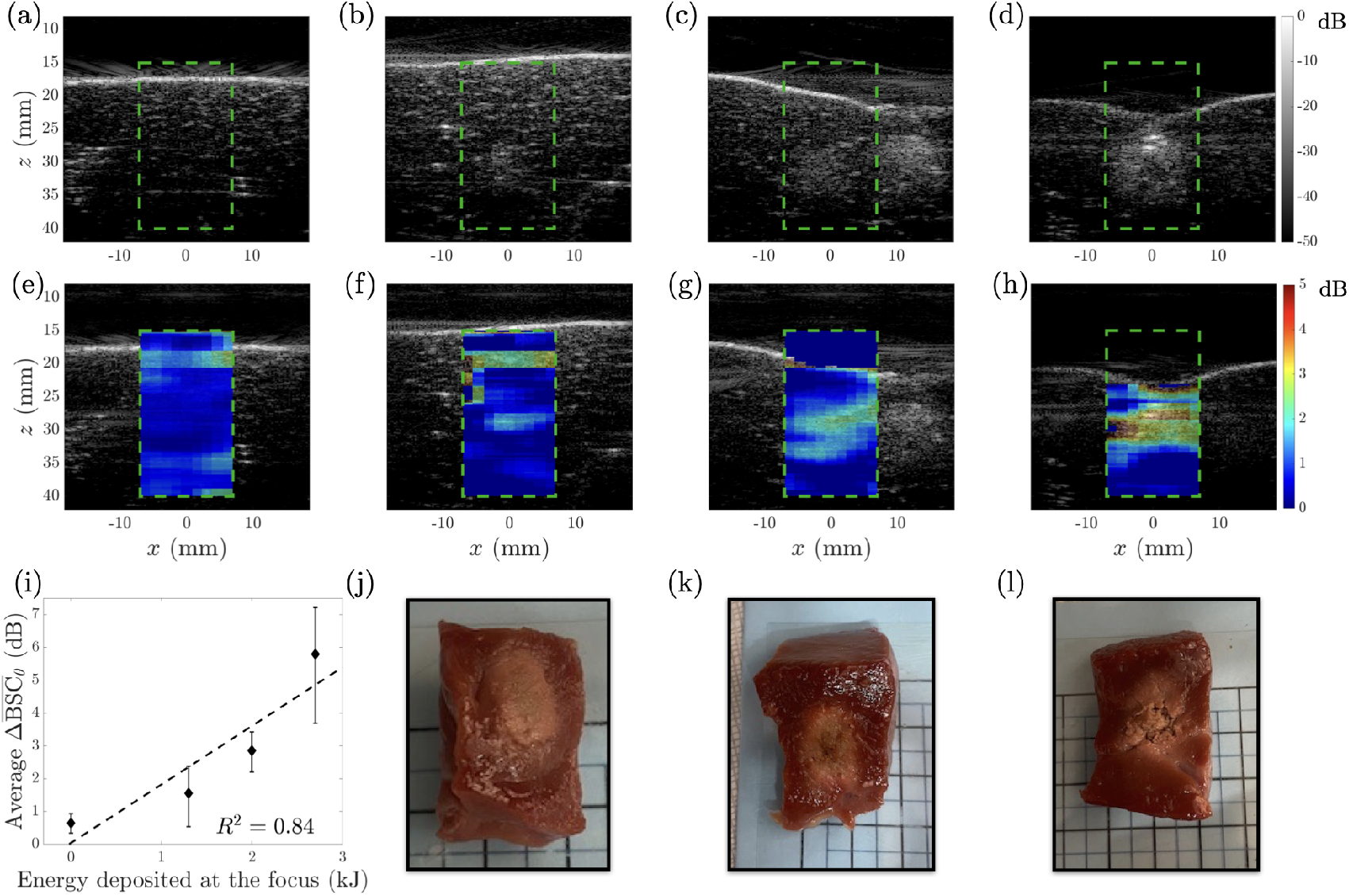
(a) B-mode image of an untreated ex vivo liver. (b), (c), and (d) B-mode images of samples 1, 2 and 3 (see Table 1). (e-h) The corresponding maps of 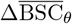 <u>for</u> (a-d) in the bandwidth range of [6-7] MHz, superimposed on the B-mode images. (i) Average value of 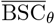 as a function of the energy deposited at the focus for samples 1 to 3 and an untreated tissue. (j), (k), and (l) Macroscopic views of samples 1, 2 and 3, respectively.

From sample 1 to sample 3, the anisotropic region progressively increases in both intensity and extent, ultimately encompassing the entire hyperechoic area visible in the B-mode image of lesion 3 (Figs. 11(d) and (h)).

## 4 Discussion

The results presented in this article demonstrate that HIFU treatments induce anisotropic alterations in the cellular architecture of liver tissue that are detectable through the angular dependance of the BSC. Histological analysis confirm the presence of anisotropy in different samples obtained during both ex vivo and in vivo treatments. The comparison with tissue heated in a water bath highlight the major role of the focused ultrasound beam in inducing the observed anisotropy within the cellular network. Histological analysis performed in the spectral domain assumes that the cellular network is responsible for the scattering. Thus, the combined analysis of the parameters *γ* and Θ makes it possible to characterize the direction and degree of alignment of the hepatocytes in the tissue. Furthermore, the agreement between the necrotic region and the anisotropic zone supports the relevance of using anisotropy as a histological biomarker for HIFU-induced tissue necrosis in the liver. The spatial and frequency scales at which these changes occur are fully compatible with clinical ultrasound imaging systems used in routine practice.

The ex vivo tissues are imaged using plane waves over a bandwidth ranging from 3.5 to 7.5 MHz, with angular apertures between ± 18°. The frequency bandwidth is chosen because it allows us to distinguish between a frequency range for which the medium appears isotropic ([3.5-5] MHz) and a range for which it appears anisotropic ([5-7.5] MHz).

A first promising correlation is found between the energy deposited at the focus and the deformation of the cellular matrix (Fig. 11(i)). This analysis will be performed in depth in future work, as it is necessary to use anisotropy as a surrogate of tissue necrosis and ultimately of the temperature reached inside the tissue. At a constant imaging frequency, the more the tissue becomes anisotropic, the smaller the angular aperture required to detect anisotropy. Therefore, the angular aperture represents a trade-off, as diffraction effects related to the imaging system limit the accessible angular range. This limitation affects the sensitivity of the measurement, since weak anisotropy can only be detected with large angular apertures. This limitation is related to the current experimental setup and does not inherently restrict the ability of the method itself to image weakly anisotropic media.

Currently, the assessment of the necrotic zone induced by HIFU is most commonly performed using MRI-T1w sequences ^36^. CT imaging is also used, offering volumetric capabilities and the advantage of being more widely accessible than MRI is. B-mode imaging is also often used to locate hyperechogenic spots induced by boiling or cavitation effects ^37^, or to evaluate devascularization when combined with the injection of US contrast agents ^38^. The adoption of multimodal strategies also represents a promising avenue to potentially eliminate the need for biopsy in specific cases. The combination of multiple imaging modalities also represents a promising field of research. For example, Apfelbeck recently investigated the combination of contrast-enhanced ultrasound imaging and multiparametric MRI^39^ in the context of prostate cancer treatment assessment.

If MRI and CT are proven to be efficient, their overall availability and cost remain significant obstacles. Furthermore, many commercial HIFU transducers induce important artifacts in MR images and show poor MRI compatibility ^40^, which encourages the development of US-based imaging methods. US-based methods are currently lacking a quantitative link between the histological features of HIFU lesions ^41^ and changes detected through imaging.

The angular and frequency dependency of scattering by soft tissue has been previously addressed in the literature through histological section analysis. Notably, the pioneering work of Mamou ^28,42^ proposed analyzing the power spectral density of three-dimensional impedance maps to identify scattering sources within soft tissues. These studies are based on a continuous distribution of impedance values, in contrast to the index ℐ introduced here, which restricts the scattering phase to the cellular network. This simplification allows a more specific investigation of the impact of the HIFU beam on this network. Attention is given only to the angular dependence of the scattering, which eliminates the need to assign an impedance value to the scattering phase.

Other important works must be mentioned here in the context of US biopathological characterization. In the recent work of Guillaumin ^43^ dealing with tumor characterization, the authors built bridges between histology and US imaging features derived from elastography and BTI ^3^. In the present article, the methodology also relies on the strong link between the microstructure and US propagation. In the future, the development of 3D imaging tools, either with a rotating 2D imaging array ^44^ or with a matrix probe ^45^, will aid in this analysis and improve the specificity of US biomarkers specificity.

## 5 Conclusion

For the first time, HIFU-induced anisotropy is observed in ex vivo and in vivo liver tissue. A first proof of concept for a new imaging method for HIFU-induced lesions based on anisotropy detection is proposed. We demonstrate that in liver tissue, BSC measurement is sensitive to the anisotropic arrangement of the cellular network. The increase in anisotropy with the amount of energy deposited in the tissue is also highlighted, paving the way for a new method to assess and monitor HIFU treatments.

## References

1 G. Cloutier, F. Destrempes, F. Yu, and A. Tang, Quantitative ultrasound imaging of soft biological tissues: a primer for radiologists and medical physicists, Insights into Imaging 12, 127 (2021).

2 F. Termite, L. Galasso, G. Capece, F. Messina, G. Esposto, M. E. Ainora, I. Mignini, R. Borriello, R. Vitiello, G. Maccauro, A. Gasbarrini, and M. A. Zocco, Multiparametric Ultrasound in the Differential Diagnosis of Soft Tissue Tumors: A Comprehensive Review, Biomedicines 13, 1786 (2025).

3 C. Papadacci, M. Tanter, M. Pernot, and M. Fink, Ultrasound backscatter tensor imaging (BTI): analysis of the spatial coherence of ultrasonic speckle in anisotropic soft tissues, IEEE Transactions on Ultrasonics, Ferroelectrics, and Frequency Control 61, 986–996 (2014).

4 C. Papadacci, V. Finel, J. Provost, O. Villemain, P. Bruneval, J.-L. Gennisson, M. Tanter, M. Fink, and M. Pernot, Imaging the dynamics of cardiac fiber orientation in vivo using 3D Ultrasound Backscatter Tensor Imaging, Scientific Reports 7, 830 (2017).

5 J.-B. Guillaumin, L. Djerroudi, J.-F. Aubry, A. Tardivon, A. Dizeux, M. Tanter, A. Vincent-Salomon, and B. Berthon, Biopathologic Characterization and Grade Assessment of Breast Cancer With 3-D Multiparametric Ultrasound Combining Shear Wave Elastography and Backscatter Tensor Imaging, Ultrasound in Medicine & Biology 50, 474–483 (2024).

6 M. F. Insana, T. J. Hall, and J. L. Fishback, IDENTIFYING ACOUSTIC SCATI’ERING SOURCES IN NORMAL RENAL PARENCHYMA FROM THE ANISOTROPY IN ACOUSTIC PROPERTIES, Ultrasound in Medicine and Biology 17 (1991).

7 B. Jiang, F. Liu, H. Fu, and J. Mao, Advances in imaging techniques to assess kidney fibrosis, Renal Failure 45 (2023), Publisher: Informa UK Limited.

8 J. Garcia-Duitama, B. Chayer, A. Han, D. Garcia, M. L. Oelze, and G. Cloutier, Experimental Application of Ultrafast Imaging to Spectral Tissue Characterization, Ultrasound in Medicine & Biology 41, 2506–2519 (2015).

9 H.-H.-P. Ngo, T. Poulard, J. Brum, and J.-L. Gennisson, Anisotropy in ultrasound shear wave elastography: An add-on to muscles characterization, Frontiers in Physiology 13 (2022), Publisher: Frontiers Media SA.

10 Q. W. Guerrero, I. M. Rosado-Mendez, L. C. Drehfal, H. Feltovich, and T. J. Hall, Quantifying Backscatter Anisotropy Using the Reference Phantom Method, IEEE Transactions on Ultrasonics, Ferroelectrics, and Frequency Control 64, 1063–1077 (2017).

11 A. P. Santoso, I. Rosado-Mendez, Q. W. Guerrero, and T. J. Hall, A Geometric Model of Ultrasound Backscatter to Describe Microstructural Anisotropy of Tissue, Ultrasonic Imaging, 016173462311711 (2023).

12 E. Franceschini, R. d. Monchy, and J. Mamou, Quantitative Characterization of Tissue Microstructure in Concentrated Cell Pellet Biophantoms Based on the Structure Factor Model, IEEE Transactions on Ultrasonics, Ferroelectrics, and Frequency Control 63, 1321–1334 (2016).

13 S. Salles, H. Liebgott, O. Basset, C. Cachard, D. Vray, and R. Lavarello, Experimental evaluation of spectral-based quantitative ultrasound imaging using plane wave compounding, IEEE Transactions on Ultrasonics, Ferroelectrics, and Frequency Control 61, 1824–1834 (2014).

14 R. O. Illing, J. E. Kennedy, F. Wu, G. R. Ter Haar, A. S. Protheroe, P. J. Friend, F. V. Gleeson, D. W. Cranston, R. R. Phillips, and M. R. Middleton, The safety and feasibility of extracorporeal high-intensity focused ultrasound (HIFU) for the treatment of liver and kidney tumours in a Western population, British Journal of Cancer 93, 890–895 (2005).

15 G. Orgera, L. Monfardini, P. Della Vigna, L. Zhang, G. Bonomo, P. Arnone, M. Padrenostro, and F. Orsi, High-intensity focused ultrasound (HIFU) in patients with solid malignancies: evaluation of feasibility, local tumour response and clinical results, La radiologia medica 116, 734–748 (2011).

16 S. Cambronero, A. Dupré, C. Mastier, and D. Melodelima, Non-invasive High-Intensity Focused Ultrasound Treatment of Liver Tissues in an In Vivo Porcine Model: Fast, Large and Safe Ablations Using a Toroidal Transducer, Ultrasound in Medicine & Biology 49, 212–224 (2023).

17 R. S. Puijk, A. H. Ruarus, H. J. Scheffer, L. G. Vroomen, A. A. Van Tilborg, J. J. De Vries, F. H. Berger, P. M. Van Den Tol, and M. R. Meijerink, Percutaneous Liver Tumour Ablation: Image Guidance, Endpoint Assessment, and Quality Control, Canadian Association of Radiologists Journal 69, 51–62 (2018).

18 V. Barrere, D. Melodelima, S. Catheline, and B. Giammarinaro, Imaging of Thermal Effects during High-Intensity Ultrasound Treatment in Liver by Passive Elastography: A Preliminary Feasibility in Vitro Study, Ultrasound in Medicine & Biology 46, 1968–1977 (2020).

19 T. Payen, S. Crouzet, N. Guillen, Y. Chen, J.-Y. Chapelon, C. Lafon, and S. Catheline, Passive Elastography for Clinical HIFU Lesion Detection, IEEE Transactions on Medical Imaging 43, 1594–1604 (2024).

20 N. T. Sanghvi, W.-H. Chen, R. Carlson, C. Weis, R. Seip, T. Uchida, and M. Marberger, Clinical validation of real-time tissue change monitoring during prostate tissue ablation with high intensity focused ultrasound, Journal of Therapeutic Ultrasound 5, 24 (2017).

21 A. Rohfritsch, V. Barrere, L. Estienne, and D. Melodelima, 2D ultrasound thermometry during thermal ablation with high-intensity focused ultrasound, Ultrasonics 142, 107372 (2024).

22 S. Zhang, S. Shang, Y. Han, C. Gu, S. Wu, S. Liu, G. Niu, A. Bouakaz, and M. Wan, Ex Vivo and In Vivo Monitoring and Characterization of Thermal Lesions by High-Intensity Focused Ultrasound and Microwave Ablation Using Ultrasonic Nakagami Imaging, IEEE Transactions on Medical Imaging 37, 1701–1710 (2018).

23 M. Monfared, H. Behnam, P. Rangraz, and J. Tavakkoli, High-intensity focused ultrasound thermal lesion detection using entropy imaging of ultrasound radio frequency signal time series, Journal of Medical Ultrasound 26, 24 (2018).

24 H. Chen, G. Y. Hou, Y. Han, T. Payen, and C. F. Palermo, Harmonic Motion Imaging for Abdominal Tumor Detection and High-intensity Focused Ultrasound Ablation Monitoring: A Feasibility Study in a Transgenic Mouse Model of Pancreatic Cancer, IEEE Transactions on Ultrasonics, Ferroelectrics and Frequency Control 62, 1662–1673 (2016).

25 G. Ghoshal, J. P. Kemmerer, C. Karunakaran, R. J. Miller, and M. L. Oelze, Quantitative Ultrasound for Monitoring High-Intensity Focused Ultrasound Treatment In Vivo, IEEE Transactions on Ultrasonics, Ferroelectrics, and Frequency Control 63, 1234–1242 (2016).

26 A. Rohfritsch, E. Franceschini, A. Dupré, and D. Melodelima, Quantitative ultrasound techniques for assessing thermal ablation: Measurement of the backscatter coefficient from ex vivo human liver, Medical Physics 50, 6908–6919 (2023).

27 Y. Fujii, N. Taniguchi, K. Itoh, K. Shigeta, Y. Wang, J.-W. Tsao, K. Kumasaki, and T. Itoh, A New Method for Attenuation Coefficient Measurement in the Liver: Comparison With the Spectral Shift Central Frequency Method, Journal of Ultrasound in Medicine 21, 783–788 (2002), Publisher: Wiley.

28 J. Mamou, M. L. Oelze, W. D. O’Brien, and J. F. Zachary, Identifying ultrasonic scattering sites from three-dimensional impedance maps, The Journal of the Acoustical Society of America 117, 413–423 (2005).

29 C. E. Zachary and S. Torquato, Hyperuniformity in point patterns and two-phase random heterogeneous media, Journal of Statistical Mechanics: Theory and Experiment 2009, P12015 (2009), 0910.2172 [cond-mat].

30 D. Garcia, L. L. Tarnec, S. Muth, E. Montagnon, J. Poree, and G. Cloutier, Stolt’s f-k migration for plane wave ultrasound imaging, IEEE Transactions on Ultrasonics, Ferroelectrics, and Frequency Control 60, 1853–1867 (2013).

31 L. X. Yao, J. A. Zagzebski, and E. L. Madsen, Backscatter coefficient measurements using a reference phantom to extract depth-dependent instrumentation factors, Ultrasonic imaging 12, 58–70 (1990).

32 J. Mamou and M. L. Oelze, Quantitative Ultrasound in Soft Tissues, volume 1, Springer Dordrecht, 2013.

33 J. J. Faran, Sound Scattering by Solid Cylinders and Spheres, The Journal of the Acoustical Society of America 23, 405–418 (1951).

34 M. S. Wertheim, Exact Solution of the Percus-Yevick Integral Equation for Hard Spheres, Phys. Rev. Lett. 10, 321–323 (1963).

35 N. W. Ashcroft and J. Lekner, Structure and Resistivity of Liquid Metals, Physical Review 145, 83–90 (1966).

36 M. Lafond, A. Payne, and C. Lafon, Therapeutic ultrasound transducer technology and monitoring techniques: a review with clinical examples, International Journal of Hyperthermia 41, 2389288 (2024).

37 L. Hong, W. Zhang, F. Pan, G. Xiaobo, H. Huang, Y. You, L. Deng, Z. Wang, and C. Zhang, An in vitro and in vivo study on extracorporeal transducer optimization for high-intensity focused ultrasound to improve the safety and efficacy of breast tumor ablation, International Journal of Hyperthermia 40, 2251734 (2023).

38 O. Rouvière, A. Gelet, S. Crouzet, and J.-Y. Chapelon, Prostate focused ultrasound focal therapy—imaging for the future, Nature Reviews Clinical Oncology 9, 721–727 (2012).

39 M. Apfelbeck, M. Chaloupka, B. Schlenker, C. Stief, and D.-A. Clevert, Follow-up after focal therapy of the prostate with high intensity focused ultrasound (HIFU) using contrast enhanced ultrasound (CEUS) in combination with MRI image fusion, Clinical Hemorheology and Microcirculation 73, 135–143 (2019).

40 N. Evripidou, A. Antoniou, L. Georgiou, C. Ioannides, K. Spanoudes, and C. Damianou, MRI compatibility testing of commercial high intensity focused ultrasound transducers, Physica Medica 117, 103194 (2024).

41 I. Rivens, C. Jayadewa, P. Mouratidis, and G. Ter Haar, Histological characterization of HIFU lesions, International Journal of Hyperthermia 41, 2389292 (2024).

42 J. Mamou, M. L. Oelze, W. D. O’Brien, and J. F. Zachary, Extended three-dimensional impedance map methods for identifying ultrasonic scattering sites, The Journal of the Acoustical Society of America 123, 1195–1208 (2008).

43 J.-B. Guillaumin, A. Nadjem, L. Vigouroux, A. Sibleyras, M. Tanter, J.-F. Aubry, and B. Berthon, 3D multiparametric ultrasound of spontaneous murine tumors for non-invasive tumor characterization, Physics in Medicine & Biology 70, 095006 (2025).

44 J.-B. Guillaumin, L. Djerroudi, J.-F. Aubry, A. Tardivon, M. Tanter, A. Vincent-Salomon, and B. Berthon, Proof of Concept of 3-D Backscatter Tensor Imaging Tomography for Non-invasive Assessment of Human Breast Cancer Collagen Organization, Ultrasound in Medicine & Biology 48, 1867–1878 (2022).

45 V. Mazellier, F. Varray, and P. Muleki-Seya, Volumetric Estimation of the Backscatter Coefficient With a Matrix Probe, IEEE Open Journal of Ultrasonics, Ferroelectrics, and Frequency Control 5, 119–122 (2025).

